# The coevolution of mammae number and litter size

**DOI:** 10.1101/2020.10.08.331983

**Authors:** Thomas A. Stewart, Ihna Yoo, Nathan S. Upham

## Abstract

Mammals are unique in provisioning their offspring with milk, lactiferous nourishment produced in glandular organs called mammae. Mammae number is hypothesized to coevolve with litter size, acting as a constraint on offspring survival. However, predicted canonical relations between mammae number and litter size (*i.e*., the ‘one-half’ and ‘identity’ rules) are untested across Mammalia. Here we analyze data for 2,301 species and show how these characters coevolve. In Mammalia, mammae number approximates the maximum reported litter size of a species, and mammae number explains more variation in litter size than other species-level traits (mass, gestation length, diet, and seasonality of contemporary geographic distribution). Clades show differences in these patterns, indicating that certain life history strategies might break the ‘rules’ of mammary evolution. Mammae number is an underappreciated constraint on fecundity that has influenced the radiation of mammals.

## Introduction

It has long been proposed that the mammae number of a species should correspond to its litter size (*1-6*). This concept is first attributed to Aristotle, who observed that species with few offspring tend to have few mammae, while species with many offspring tend to have many mammae (*1*). Previous interspecific analyses identified a linear relationship between these traits, where the average number of offspring per litter is approximately one-half of mammae number (*7, 8*). This ‘one-half rule’ is predicted to be general for mammals (*9*), and species that deviate from this pattern are considered exceptional (*10*). It has been further noted that the maximum reported litter size of a species approximates mammae number (*7*), herein referred to as the ‘identity rule.’ As a corollary, it is hypothesized that mammae number constrains litter-size evolution by limiting the number of offspring that can have access to milk and survive (*7, 11*). However, it is unclear whether the particular numerical relation between mean litter size and mammae number reflects their coevolution directly, or whether it might instead be an indirect consequence of the coevolution between mammae number and maximum litter size.

General models of mammary evolution for Mammalia are challenged by several observations. First, lineages show remarkable diversity in their life histories and habits (*12*). For example, patterns of fetal *versus* post-birth maternal investment can differ dramatically between clades (e.g., marsupials and placentals (*13, 14*)). In some lineages, mothers will nurse offspring from multiple pregnancies concurrently (e.g., humans and macropod marsupials (*13*)). And females can nurse offspring that are not their own (*i.e*., allonursing (*15*)). Second, the studies that proposed the one-half and identity rules had limited taxonomic sampling to address mammal-wide patterns (90 placental species (*8*), and 266 rodent species (*7*)), and they did not apply phylogenetic comparative methods, which are critical for evolutionary interpretations of character correlation (*16*). Finally, several other ecological and physiological traits are known to correlate with litter size in mammals (*17-22*), raising questions about the explanatory power of the one-half and identity rules. Here we survey mammalian species diversity broadly and apply phylogenetic tests to ask (i) whether and how mammae number coevolves with litter size across Mammalia and (ii) how mammae number compares to other known predictors of litter size. We show that mammae number and fecundity are fundamentally linked and argue that mammae number likely constrains litter-size evolution.

## Results

### Testing the one-half and identity rules

Data on mammae number, mean litter size, and maximum litter size were collected from the literature and museum specimens for 2,301 species (Fig. 1a, Supplementary Table 1) and mapped to a time-calibrated molecular phylogeny of extant mammals (Fig. 1b, Supplementary Fig. 1) (*23*). Major clades differ in their patterns of variation of these characters (Fig. 1b, Supplementary Fig. 2, Supplementary Table 2). Mammae number is typically even, with nipples positioned bilaterally along the milk lines; however, some marsupials have an odd number of mammae and nipples on the midline (*e.g., Monodelphis domestica*, Fig. 1c) (*13*). Lineages also differ in their distributions of litter-size values (Fig. 1b; Supplementary Fig. 2b,c; Supplementary Table 2). For instance, chiropterans (bats) and primates tend to have fewer offspring than rodents or carnivorans (cats, dogs, and their allies). Notably, the afrotherian *Tenrec ecaudatus* is the mammal with the greatest number of mammae, the largest mean litter size, and it is tied with the opossum *Didelphis virginiana* for the largest maximum litter size (Fig. 1c).

**Fig. 1.**
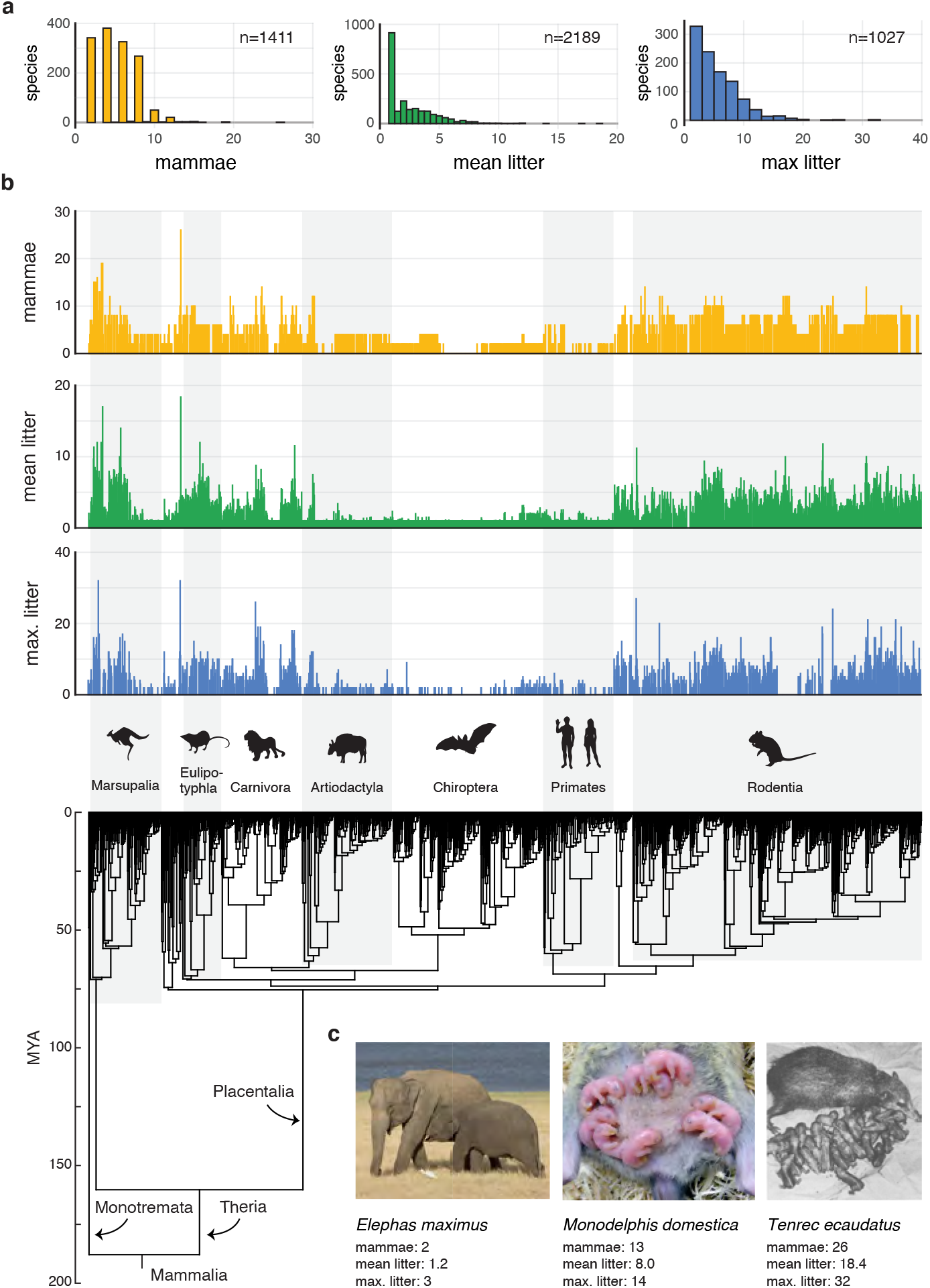
Mammals show dramatic diversity in mammae number and litter size. (a) Histograms of available data on mammae number, mean litter size, and maximum litter size, (b) Characters plotted on a species-level molecular phylogeny of Mammalia (25) pruned to the 2,301 branches with at least one data type. (c) Photographs of mammals highlight their diversity in reproductive strategies: Mother *E. maximus* suckling one calf from a pair of mammae positioned between the forelimbs (photo: Carlos Delgado with permission). Mother *M. domestica* suckling its pups from an odd number of inguinally positioned mammae (photo: Wheaton et al. (26) with permission). Mother *T. ecaudatus* and 31 offspring born of a single litter (photo: Louwman(27) with permission).

The one-half rule predicts that a linear regression of mean litter size ~ mammae number should recover a slope of 0.5 and intercept of zero, and the identity rule predicts that a linear regression of maximum litter size ~ mammae number should recover a slope of 1.0 and intercept of zero (*7*). To test these hypotheses, we conducted phylogenetic generalized least squares (PGLS) regression (*24*), accounting for topological and age uncertainty using a sample of 100 trees drawn from the node-dated credible tree set (*23*). In Mammalia, mammae number coevolves with mean litter size (*P* values < 0.05 across all 100 trees, median R^2^: 0.28). However, this relation does not follow the one-half rule: across 100 trees, the PGLS slope mean is 0.44, and 95% confidence intervals (CIs) do not contain 0.5; the PGLS intercept mean is 0.65 and 95% CIs do contain zero (Fig. 2a, Supplementary Table 3). PGLS regressions also show that mammae number coevolves with maximum reported litter size in Mammalia (*P* values < 0.05 across all 100 trees, median R^2^: 0.30). This relation does follow the hypothesized identity rule: across 100 trees, the slope mean is 0.97, intercept mean is 0.54, and 95% CIs of both parameters contain the predicted values of one and zero, respectively (Fig. 2b, Supplementary Table 3).

**Fig. 2.**
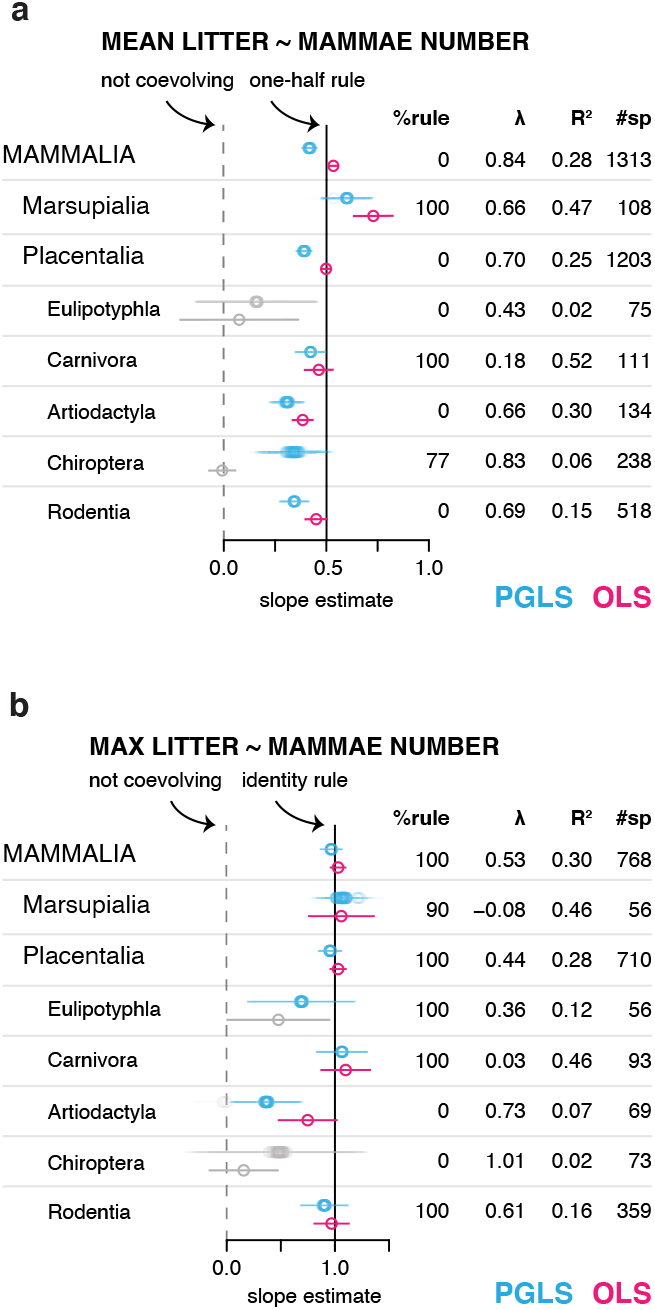
Mammae number coevolves with litter size. Phylogenetically generalized least squares (PGLS) and ordinary least squares (OLS) regressions were run for (a) mean litter size ~ mammae number and (b) maximum litter size ~ mammae number. PGLS analyses were run on 100 trees to account for phylogenetic uncertainty. OLS regressions do not take the phylogenetic relationships of species into account and are presented to contextualize PGLS regressions. Slope estimates are indicated with a circle, and 95% confidence intervals (CIs) of each estimate are shown with horizontal lines. If PGLS or OLS regressions hadP value > 0.05, the estimate is plotted in grey. In the table to the right, ‘%rule’ is the percentage of PGLS regressions that were significant at P 0.05 and for which the slope and intercept 95% CIs contained the predicted values. Other columns show the mean Pagel’s λ for the 100 trees (a measure of phylogenetic signal), the median variance explained (R^2^), and the total number of species in the clade that were analyzed in the regression. PGLS and OLS regressions are further summarized in Supplementary Tables 3,5.

Not all clades follow the identity rule. In Chiroptera, these traits are not coevolving (PGLS regressions have *P* value > 0.05 across all 100 trees) (Fig. 2b). This result might reflect some bats having evolved non-milk producing mammae, which are used by offspring for attachment during flight (*12*). In Artiodactyla, mammae number and maximum litter size are weakly coevolving (PGLS regressions have *P* value < 0.05 across 99 trees, median R^2^: 0.07) but with a shallower slope than predicted by the identity rule (Fig. 2b). This divergence from the general mammalian pattern might reflect the tendency of ungulates to give birth to highly developed, precocial offspring and the presence of udders in some species.

### Relative predictive strength of mammae

Litter size is known to correlate with a number of other traits besides mammae number, including adult body mass (*17*), gestation length (*18*), dietary category (*19*), and environmental features (*e.g*., latitude (*17*), habitat type (*20*), geographic region (*21*), and climatic seasonality. (*22*)). To analyze the explanatory power of mammae number on litter-size evolution relative to ecological and life-history characters, we constructed an additive PGLS model and measured the contribution of each predictor to the model’s total fit over 100 trees. Five variables were selected from a broader set of nine to minimize issues of multicollinearity in the model and to maximize phylogenetic coverage (Supplementary Fig. 3): mammae number, adult body mass, gestation length, amount of plant-based matter in the diet, and the standard deviation of quarterly temperatures over the contemporary distribution of each species as a metric of climatic seasonality (see Methods for sources of predictor variable data).

In Mammalia, mammae number predicts both mean and maximum litter size more strongly than any other variable analyzed (Fig. 3a; Supplementary Figs. 4, 5; Supplementary Table 4). Mammae number produces the greatest ΔAIC in 99% and 98% of trees when mean and maximum litter size, respectively, are considered (Fig. 3b). This finding is unexpected because the substantial literature on mammalian litter-size evolution rarely discusses mammae number as a predictor. More often, the energetics of lactation and body mass scaling are highlighted as determining litter size variation (*e.g*., refs. (*17, 25*).

**Fig. 3.**
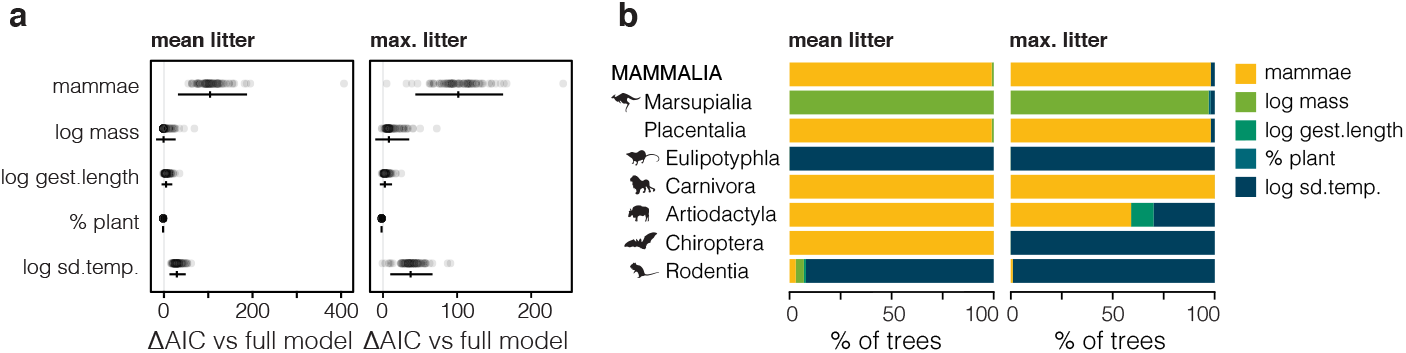
Mammae number predicts litter size more strongly than other traits. Likelihood testing was used to compare a five-predictor model with models that sequentially excluded one of the predictor variables: mammae number, log adult body mass, log gestation length in days, percentage of plant-based matter in the diet, and seasonality of contemporary geographic distribution of each species (calculated by log of the standard deviation of quarterly temperatures in degrees C). Analyses were run with either mean or maximum litter size as the response variable across 100 trees to account for phylogenetic uncertainty, (a) Differences in Akaike Information Criterion (ΔAIC) metrics were calculated between the five-predictor model and each of the four-predictor models. In Mammalia, mammae number consistently yields the greatest ΔAIC. (b) Stacked bar graph showing the predictor variables that yielded the greatest ΔAIC in the 100 trees. Analyses were run for Mammalia and clades where data were available for at least 40 species.

Clades differ in which predictor variable produces the greatest ΔAIC (Fig. 3b, Supplementary Fig. 3, 4, Supplementary Table 4). Thus, the relative explanatory power of mammae number on litter size evolution is heterogeneous across Mammalia. This disparity appears to reflect lineages having distinct ecological and life-history strategies. For example, in rodents, seasonality predicts maximum litter size more strongly than mammae number (species that occupy more seasonal environments have larger litter sizes than species in less seasonal environments; Supplementary Fig. 5c), while in carnivores, mammae number is the most strongly predictive variable. Both Rodentia and Carnivora evolve according to the identity rule (Fig. 2b), however differences in their geographic distributions or relation to the environment (e.g., length of breeding seasons or predictability of food availability) determine the strength of the correspondence between mammae number and litter size.

## Discussion

Overall, these analyses both confirm and add novel detail to the long-standing prediction that mammae number and litter size are fundamentally related (*1-8*). We find support for the hypothesized ‘identity rule’ across Mammalia, where mammae number evolves with a one-to-one correspondence to the maximum observed litter size of a species. Maximum values analyzed here can be understood as low frequency litters of substantially greater than average magnitude and not the physiological limits of species. Thus, consistent with previous predictions (*7, 8*), mammae number evolves to provide a factor of safety for rare large-litter events. If mammae number becomes canalized in a lineage, it will affect their ability to evolve larger litter sizes by establishing an anatomical limit to a mother’s nursing capacity. The specific numerical relations between mammae number and mean litter size (e.g., PGLS slope mean of 0.44 in Mammalia) reveals that safety factors are generally greater than what was predicted by the one-half rule.

Clades can deviate from this general mode of mammary evolution and show disparity in the explanatory power of mammae number on litter size. Residuals of the PGLS regression of maximum litter size ~ mammae number point toward how taxa achieve this divergence. The most extreme case is the naked mole-rat, *Heterocephalus glaber*, which has the largest residual among mammals (Supplementary Fig. 6) and the most highly derived eusocial habits of any mammal, with all colony members supporting reproduction of a single breeding female (*10*). Discovering how certain behaviors (e.g., mammae-sharing among litter mates (*14*), tenacious suckling (*11*), allonursing and other types of cooperative breeding (*15*)), as well as the ecological and life-history strategies of species, contribute to this disparity should be the focus of future research. Investigations into the developmental basis of mammary patterning, how mammae number becomes canalized, and how supernumerary structures are transformed into functional mammae are also necessary. Ultimately, understanding the remarkable variation in mammae number across Mammalia will depend upon integrating causal explanations of both general patterns and deviations from evolutionary ‘rules’ at shallower phylogenetic scales.

## Materials and methods

### Collection of mammary and litter data

Data on mammae number and litter size were compiled from the literature, online databases, and the in person study of museum specimens. In total, 12,994 unique records were collected. Major data sources include a compendium of mammalian reproductive biology (*12*), personal correspondence with Dr. Avery Gilbert, who shared his original notes and data from a previous study of the one-half rule in rodents (*7*), the museum specimen database VertNet (*28*), and the trait compendium PanTHERIA (*29*).

Over the years, more than 250,000 unique taxonomic identifiers (genus, species, and subspecies name combinations) have been applied to the approximately 6,000 currently recognized extant mammalian species (*23, 30*). To account for changes in taxonomy, taxonomic identifiers in raw datasets were matched with a synonymy table by Meyer and colleagues (*30*). Raw data identifiers that could not be matched to the table were investigated manually to diagnose potential misspellings or taxonomic changes not captured by the table. In total, data were collected for 2,698 mammal species belonging to 1,016 genera, 152 families, and 27 orders (of 1,314 genera, 167 families, and 27 orders included in the latest mammal checklist (*31*)). The number of records for a species ranged from 1 to 113. If multiple records of a character were collected for a species, then the records were collapsed so that each species was represented by a single estimate of mammae number, mean litter size and maximum litter size.

Mammae number was collected for 1,645 species. In monotremes, mothers suckle offspring from a pair of milk patches in the pouch (*32*) and are, thus, coded as having 2 mammae. In therians, the clade comprised of marsupials and placentals, mothers suckle offspring from nipples or teats, and mammae number was determined by nipple or teat count (*33*). Some mammals have mammae that are not used for suckling (*e.g*., the pelvic teat pair of some bats (*12, 34*) and the anterior nipples of Shar Pei dogs (*35*)). We report the counts of all mammae, not only those that produce milk, because information on the milk production of individual mammae is unavailable for most species. In the compendium of mammalian reproductive biology by Asdell (updated by Hayssen et al. (*12*)), mammae counts are sometimes reported at the genus level; these values were extrapolated to the species level only for species that are named explicitly in the associated text.

Generally, only a single mammae value is reported for a species. This is interpreted to be the modal value for the species. Within a species, mammae number can vary. For example, human mammae counts can vary from zero (*36*) to ten (*37*), and *Tenrec ecaudatus* can vary from 14 to 34 (based upon measurements by Dr. Link Olson of 26 specimens at the Field Museum of Natural History (Chicago, USA) and the Université d’Antananarivo, Département de Biologie Animale (Antananarivo, Madagascar)). If a reference reported a range of mammae values for a species *(e.g., Alopex lagopus* with 12-14 mammae (*38*)) but did not specify a typical, or modal value, then the reference was excluded from the dataset. If multiple references reported different mammae count for a particular taxon, as was the case for 91 species in our analyses, then we prioritized one reference as the modal value. When deciding which value to designate as the modal estimate, we preferentially selected reports from the primary literature over online databases and textbooks. In several lineages, mammae number differs between mates and females (*e.g*., horses) (*39, 40*). In such cases, we report and analyze the mammae number of females. All references to odd numbers of mammae were investigated manually to check for errors in data entry.

Mean litter size data were collected for 2,498 species, and maximum litter size estimates were collected for 1,118 species. To estimate litter size, we considered only direct observations of offspring number. Indirect anatomical proxies of litter size (e.g., placental scars *(41))* were not included in the analyses. Both pre-birth and post-birth counts of offspring number are included in our data. Pre-birth observations differed slightly among clades: Monotremes are oviparous, and therefore estimates of litter size included observations of embryos still in the eggshell. Marsupials have short periods of embryonic implantation; therefore, estimates of litter size in marsupials considered observations of late-stage embryos not yet implanted and implanted embryos. Therians have comparatively longer pregnancies, and so estimates of litter size included implanted embryos. When available, the timepoint at which the litter size was measured was recorded. We analyzed how the time point of sampling could impact our analyses by re-running analyses on datasets partitioned to include only pre-birth or only post-birth observations (discussed below). Collectively, data on litter size comprise approximately 497,000 individual litters, as estimated by the sum of all litters observed to generate each species’ litter estimate.

Data on the litter size of a particular species was frequently available from multiple sources. If multiple reports of litter size were available, they were combined to generate a single value for each species. To consolidate the multiple mean litter size values of a species, we calculated a weighted mean, accounting for the number of individual litters observed per study (*e.g*., if study A reports mean litter of 1.5 by observing 5 litters and study B reports mean litter of 1.3 observing 15 litters, then a weighted mean of 1.35 was used in our analysis). Publications that reported a mean litter size but did not report how many individual litters were sampled were treated as a single observation. To consolidate multiple records of maximum litter size, we selected the greatest value reported of all records available.

In our dataset, humans are reported as having a maximum of three children per pregnancy. This value is less than the maximum number of offspring ever reported for a human pregnancy (eight children surviving (*42*)), but was selected based on two criteria. First, the highest reported number of children in a single human pregnancy likely reflects fertility treatments and *in vitro* fertilization techniques, not a number typical of humans. Second, estimates of maximum litter size will be impacted by the number of pregnancies sampled. In humans, triplet rates are approximately 1/1000 births (*43*), and a sample of 1,000 observed pregnancies is consistent with the general upper limits of the number of sampled litters for other species in the dataset. Therefore, we selected the maximum offspring estimate that would likely have been observed if human pregnancies were sampled at the same rate as other species in the dataset.

The litter size data published by PanTHERIA includes values that were transformed according to a model weighting central tendency of multiple variables at the ordinal level (*29*). As a result, some litter size values in the PanTHERIA database are less than 1.0, despite it being a biologically meaningful minimum threshold of litter size. In our dataset, there are 21 such species (values range from 0.96 to 0.99). We chose to include these species, because the approximately 0.04% error introduced by this transformation is likely comparable to error rates of other taxa with small sample sizes of observed litters.

Character data includes only extant species. Fossilized mammary tissues have not been described(*44*). Therefore, fossil taxa cannot be included in the analyses presented (PGLS regressions or likelihood modeling). Fossilized embryos have been described for several crown-group therians (*45*) (*i.e*., bats (*46*), cetaceans (*47*) (although see (*48*)), equids (*49*), oreodonts (*50*), mammoth (*51*) and ground sloths (52)). In all cases, either one or two embryos are preserved. These values are consistent with mean litter size values among extant representatives of these clades, which tend to have few and relatively large precocial young.

### Descriptive statistics of reproductive characters

We tested whether clades differed from one another in the distribution of mammae number, mean litter, and maximum litter values. We selected the following clades for their large sample sizes and suitability for later analyses: Marsupials, Eulipotyphla, Carnivora, Artiodactyla, Chiroptera, Primates, and Rodentia.

First, we tested whether the data were significantly different from the normal distribution using the Shapiro-Wilk normality test (*53*). The distributions of all three variables differed from normal (mammae number: W = 0.89711, *P* value < 2.2e-16: mean litter: W = 0.79962, *P* value < 2.2e-16; maximum litter: W = 0.84794, *P* value 2.2e-16). Therefore, we proceeded with non-parametric tests for differences between clades. We next tested for differences among clades, we used the Kruskal-Wallis rank sum test (*53*). For each variable, we detected differences among clades (mammae number: chi-squared = 782.25, *P* value < 2.2e-16, df = 6; mean litter: chi-squared = 1202.1, *P* value < 2.2e-16, df = 6; maximum litter: chi-squared =: 331.38, *P* value < 2.2e-16, df = 6). Finally, to identify which clades differed from one another, we applied *post hoc* pair-wise comparisons with the Tukey-Kramer test(*54*) with Tukey-Dist approximation for independent samples using the R package “PMCMR” (*55*). For each variable, clades were diagnosed as differing if *P* value for this test was < 0.05.

### Phylogeny

The phylogenetic framework for this study is the 4,098 species-level time-calibrated molecular phylogeny of mammals that was recently published by Upham and colleagues (*23*). A maximum clade credibility consensus tree of this ‘DNA-only’ data set is used to display the data in Fig. 1b and Supplementary Fig 1. To account for phylogenetic uncertainty and to improve parameter estimates (*56*), all analyses were run on 100 trees from the credible set of Mammalia phylogenies (downloaded at http://vertlife.org/data/mammals).

To test sensitivity of our results, we also analyzed mammae and litter size coevolution using the ‘completed’ trees of Upham and colleagues (*23*), which are 5,911-species phylogenies that have DNA-missing species imputed within taxonomic constraints across the credible set of trees. Potential issues can arise when trait evolution analyses are run on trees generated by taxonomic imputation (*23, 57*), and for this reason we prioritize the DNA-only dataset in the paper. Nevertheless, the more inclusive 5,911-species phylogeny allows a means to explore the robustness of our results using an expanded mammae and litter size dataset. Major results presented in the manuscript are robust to both approaches, although parameter estimates and confidence intervals (CIs) can show minor differences (compare Figs 2, 3 to Supplementary Fig 7).

### Regression analyses

Phylogenetic generalized least squares (PGLS) regressions (*24*) were conducted in R using the ‘gls’ function in the package ‘ape’(*58*). The regression models were fit with simultaneous estimates of Pagel’s λ using maximum likelihood (*59*). Ordinary least squares (OLS) regressions were conducted in R using the standard function ‘lm’ as a means of comparing non-evolutionary models to PGLS models, which account for expected covariances based on shared evolutionary history among species. Regressions were run for all Mammalia and on ordinal and infraclass clades for which data was available for at least 40 species.

We assessed whether results of the univariate PGLS analyses shown in Fig. 2 might be disproportionately impacted by outlier species in our dataset. To do this, residuals from the PGLS models run on each of 100 trees were multiplied by the factorized positive-definite matrix of phylogenetic variancc-covariancc. and then divided by the standard deviation to place in comparable units. Neither of the regressions for Mammalia have species with residuals with an absolute value equal or greater than 3 (Supplementary Fig. 8), a common threshold for defining outliers (*60*). Therefore, analyses are presented on the complete dataset (Supplementary Table 1).

### Collection of ecological and life-history trait data

To assess the relative predictive effect of various traits hypothesized to impact litter size, an additive linear model was constructed. Nine predictor variables were initially considered: (1) mammae number; (2) adult body mass (kg); (3) percentage of plant-based matter in the diet; (4) gestation length (days); (5) generation length (days); (6) weaning length (days); (7) temperature seasonality on the contemporary geographic distributions of each species; (8) mean latitude of the same contemporary geographic distribution; (9) mean temperature of the contemporary geographic distribution of each species. Data for the life-history and ecological characters were collected from a variety of sources and unified to the full 5,911-species mammal phylogeny, accounting for changes in taxonomy using the synonym table as described above (Supplementary Table 6).

Estimates of mean body mass were assembled from the databases of Faurby and Svenning (*61*) (5,351 species) and EltonTraits (*62*) (46 species, only those coded with certainty level “1” for direct observations). The mass of sexually dimorphic species was considered as a species-level mean of adult males and females (*63*). To develop a metric of trophic level, we modified the EltonTraits (*62*) diet classification into a total dietary percentage of plant-based matter by summing the herbivorous categories of Diet.Fruit, Diet.Nect, Diet. Seed, and Diet.PlantO and then dividing by 100 to yield a proportion (5,054 species). To estimate generation length, we used the calculations of Pacifici *et al*. (*64*), which are based on species reproductive life span, from the age of first to last reproduction (5,322 species). For gestation and weaning length, we extracted these variables from the Amniote database compilation (*65*) (2,136 and 1,963 species, respectively).

To collect data on abiotic variables, we generated species-level summaries by intersecting remotely sensed global layers with ~1-degree equal area grid cells (360 x 114 km at the equator), and then taking the mean of occupied grid-cell values on a per-species basis. Grid cell occupation was determined as >20% cell coverage by a species’ geographic range polygon (or, if <20% of any cell was covered, then the cell of greatest coverage) using the mammal range polygons updated from IUCN (*66*) to match the taxonomy of to the mammal phylogeny (*23, 67*). To calculate climate seasonality, we extracted annual temperature data from CRU CL 2.0 (*68*) (downloaded at https://crudata.uea.ac.uk/cru/data/hrg/tmc/), which is a gridded climatology of 1961 to 1990 monthly means at a 10-minute longitude/latitude resolution. We then aggregated those temperature values to 1-degree equal area grid cells and calculated the standard deviation of quarterly mean values per cell as a metric of annual temperature seasonality (5,782 species).

Of those nine predictor variables (Supplementary Table 6), a subset was selected according to criteria aiming to retain (i) the broadest phylogenetic sampling; (ii) overlap with our mammae and litter size dataset; and (iii) minimize issues of collinearity among the predictors. Specifically, we followed the rule of thumb by Tabachnick & Fidell (*69*) that predictor variables included in a multiple regression analysis should have a bivariate correlation of less than 0.70. All positive right-skewed variables (body size, generation, gestation, and weaning lengths, and abiotic variables) were log-transformed to reduce residual heteroscedasticity prior to analyses. Spearman’s correlations were calculated (Supplementary Fig. 3), and on the basis of these comparisons we chose to exclude generation length and weaning length for their high correlation with one another and with log. mass. Of the abiotic predictors, latitude and temperature were excluded for their high correlation with one another and with temperature seasonality (‘log. sd.temp’). Temperature seasonality was prioritized for best representing the long-standing hypothesis that shorter seasons might select for larger litters (*22*).

For the remaining five potential predictors, we additionally analyzed their variance inflation factors (VIF) to evaluate collinearity. Such approaches cannot be applied to PGLS models directly, and so a non-phylogenetic version of the five-parameter model was run in R using the standard function ‘lm.’ VIF were calculated using the R package ‘car’ and the function ‘vif on the model output. VIF values greater than 10 would suggest strong collinearity *(70).* When run for Mammalia, the maximum VIF value of the five variables is 2.50 when mean litter size is the response variable and 2.66 when maximum litter size is the response variable. Therefore, all five predictors were used in the model.

### Likelihood model testing

To calculate the relative predictive effect of each variable upon litter size (either mean or maximum), first the full five-parameter additive model was run. Then, each of the predictor variables was sequentially dropped from the model, and each of the resulting four-variable models were run. Comparisons between the full model and each of the four-parameter models were made using Akaike information criterion (AIC) and sequential sums of squares (ΔR^2^) using the ‘anova.pgls’ function in the R package ‘caper’ (*71*). These model comparisons were conducted on each of 100 trees from the credible set of Mammalia phylogenies, and summarized to determine which of the predictors explained the greatest amount of variation in litter size. For plotting, the effects of each predator variable were standardized by mean centering and standard deviation scaling using the ‘scale’ function in base R.

### Impact of timepoint of litter sampling

As noted above, estimates of litter size in our dataset contain both pre- and post-birth observations. Litter size measures can be impacted by the timepoint of pregnancy when offspring are counted, particularly in marsupials, in which it has been observed in some species that more eggs are fertilized than are expected to survive through weaning (*13, 21, 72*). To test the sensitivity of our results to the effect of sampling timepoint and the impact of offspring attrition prior to weaning, we subset data into either pre-birth or post-birth categories of observation. Most records of litter size in our dataset do not mention observation timepoint explicitly, and these records were excluded from our pre-*versus* post-birth comparisons. PGLS analyses were repeated for ordinal and superordinal clades where data were available for at least 40 species.

Results of the PGLS regression of mean litter ~ mammae number are consistent between pre- and post-birth datasets (Supplementary Fig. 9a). These partitioned data sets can produce results that differ from analyses of the full data set (*e.g*., slope estimates in Mammalia and Artiodactyla), a difference that we attribute to approximately 30% reduction in taxonomic coverage size for these partitioned datasets. PGLS regressions of maximum litter ~ mammae number show minor but consistent differences between pre-birth and post-birth datasets (Supplementary Fig. 9b). For Mammalia, Placentalia, and Rodentia, the slope mean estimate is greater in the pre-birth data set than for the post-birth data. These differences, although subtle, might reflect embryonic attrition and a tendency for the slope of relationship to reduce during the time course of pregnancy and weaning. However, this pattern is difficult to disentangle from the approximately 50% reduction in taxonomic coverage between the full and partitioned data sets. Nevertheless, the major result that in Mammalia mammae number and maximal reported litter size evolve according to the identity rule is recovered under all conditions. Marsupials, the clade where we expect this effect to be most dramatic, because some species regularly produce more embryos than have available teats (*13, 21, 72*), could not be considered in this manner, as our dataset contains few records where a range of observed embryos and where the time point of observation is specified.

### Licenses for photographs and silhouettes

The photo of the opossum in Fig. 1 is from Wheaton *et al*. (26) which is an open-access article distributed under the terms of the Creative Commons Attribution License and allows for unrestricted use, distribution, and reproduction with citation. The photo of the elephant in Fig. 1 is from Carlos Delgado and made available on Wikipedia, distributed under Terms of the Creative Commons license for Attribution-ShareAlike 4.0 International (CC BY-SA 4.0), which allows sharing of content in any format. The photo of the Tenrec in Fig. 1 is from Louwman (*27*). John Wiley and Sons publishing and its Copyright Clearance Center granted access for usage of this photograph for use.

Silhouette for the marsupial clade is by AussieIcons and licensed as Freeware, which means it is free for non-commercial use (https://www.fontspace.com/web-dog/aussieicons). Silhouette for the carnivoran clade is by Iconian Font, Zoologic and licensed as Freeware, which means it is free for non-commercial use (https://www.fontspace.com/iconian-fonts/zoologic). Silhouette for the artiodactyl clade is by Zoologic and licensed as Freeware, which means it is free for non-commercial use (https://www.fontspace.com/iconian-fonts/zoologic). Silhouette for the eulipotyphlan clade is by T Michael Keesey after Monika Betley (phylopic.org, CC-BY4; phylopic.org, CC-BY3). Silhouette for the chiropteran clade is by Yan Wong (phylopic.org, CC-BY4; phylopic.org, CC-BY3). Silhouette for the rodent clade is by Rebecca Groom (phylopic.org, CC-BY4; phylopic.org, CC-BY3). Licensing terms of the three PhyloPic silhouettes allow them to be used freely for any purpose. Silhouette of humans is by author T.A.S. and based upon an illustration from the plaque of the Pioneer spacecraft.

## Acknowledgments

Thanks to J. Miyamae, B. Aiello, D. Krentzel, G. Slater, M. Zeldich, Z.Y. Wang and N. Piland for discussion and feedback on the manuscript. Thanks to A. Gilbert for rodent data and L. Olson for Tenrec data. Funding: Research supported by The University of Chicago, the Brinson Foundation, and NSF grant DEB-1441737.

## Author contributions

TAS and NSU conceived of the project. All authors collected and curated the data. TAS and NSU conducted analyses and wrote the paper. IY helped with data visualization and edited the paper.

## Data Accessibility

Data analyzed in this paper are provided in Supplementary Tables 1 and 6.

## Code Accessibility

All code used in the analysis are freely available online at https://github.com/ThomasAStewart/mammae_project

## Supplementary Information

**Supplementary Fig. 1.**
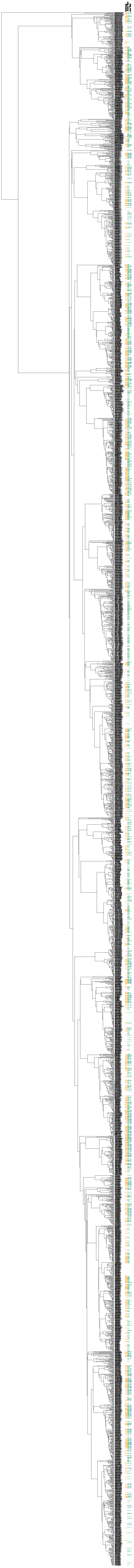
Plotting character presence/absence on a 4098 species phylogeny of mammals. Data are mapped on to the time-calibrated molecular phylogeny analyzed in the present study, which was published in 2019 by Upham and colleagues (*23*). The maximum clade credibility topology shown is from the credible set of 10,000 DNA-only node-dated trees and downloadable at http://vertlife.org/phylosubsets. Available character data for mammae number, mean litter size, and maximum litter size are denoted with colored marks next to the taxon names.

**Supplementary Fig. 2.**
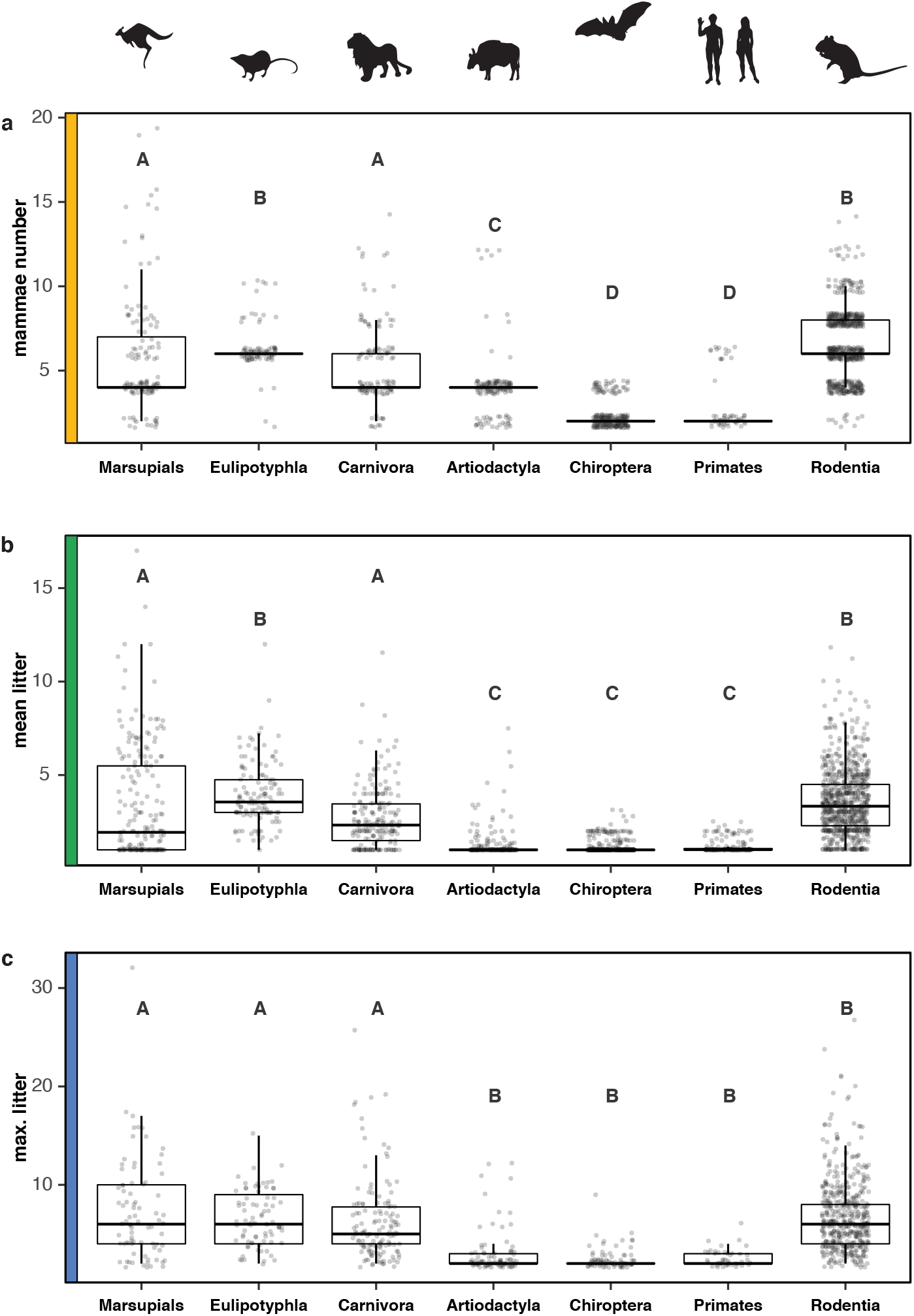
Testing for clade-wise differences in mammae number, and mean and maximum litter size. (a) Mammae number, (b) mean litter size, and (c) maximum litter size values of species are plotted for major lineages. Box and whisker plots show median values and the first and third quartiles, A Tukey-Kramer *post-hoc* test was used to detect clades that differ from one another in their distributions to a threshold *P* value’< 0.05. These analyses diagnose four significantly different groups for mammae number, three for mean litter size, and two for maximum litter size. These groups are denoted with uppercase letters A-D on the corresponding plots. Data points of mammae number are jittered both laterally and vertically to better show the distribution of overlapping values. Data points of litter size were jittered only laterally. The phylogenetic relationships among clades were not considered in this test, but divergence times all exceed 60-million years (*23*).

**Supplementary Fig. 3.**
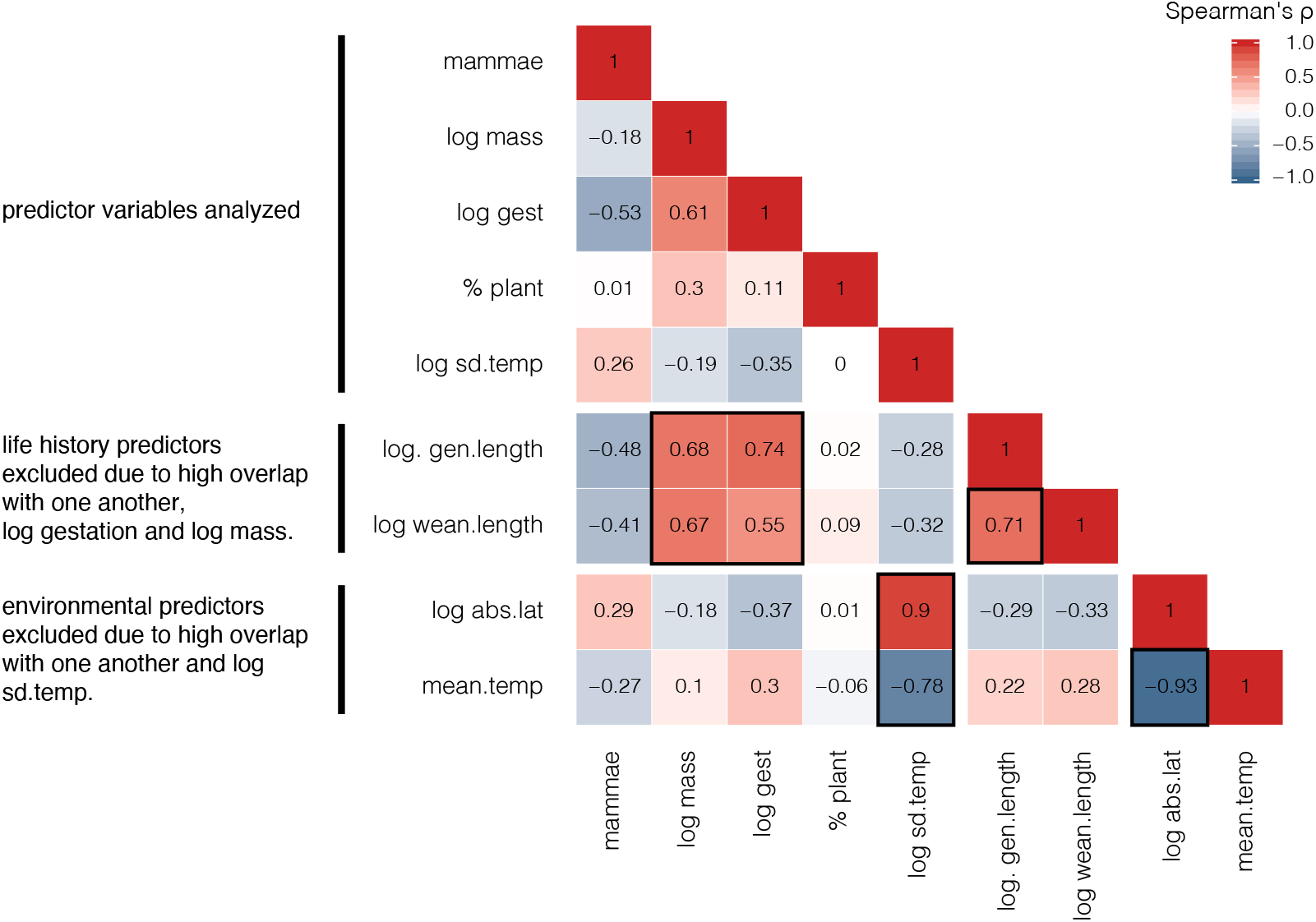
Spearman’s correlation matrix of litter-size predictor variables. Heatmap of reported values shows Spearman’s ρ for the nine predictors that were considered; five were included in the additive multivariate PGLS regression, and four were excluded from analyses.

**Supplementary Fig. 4.**
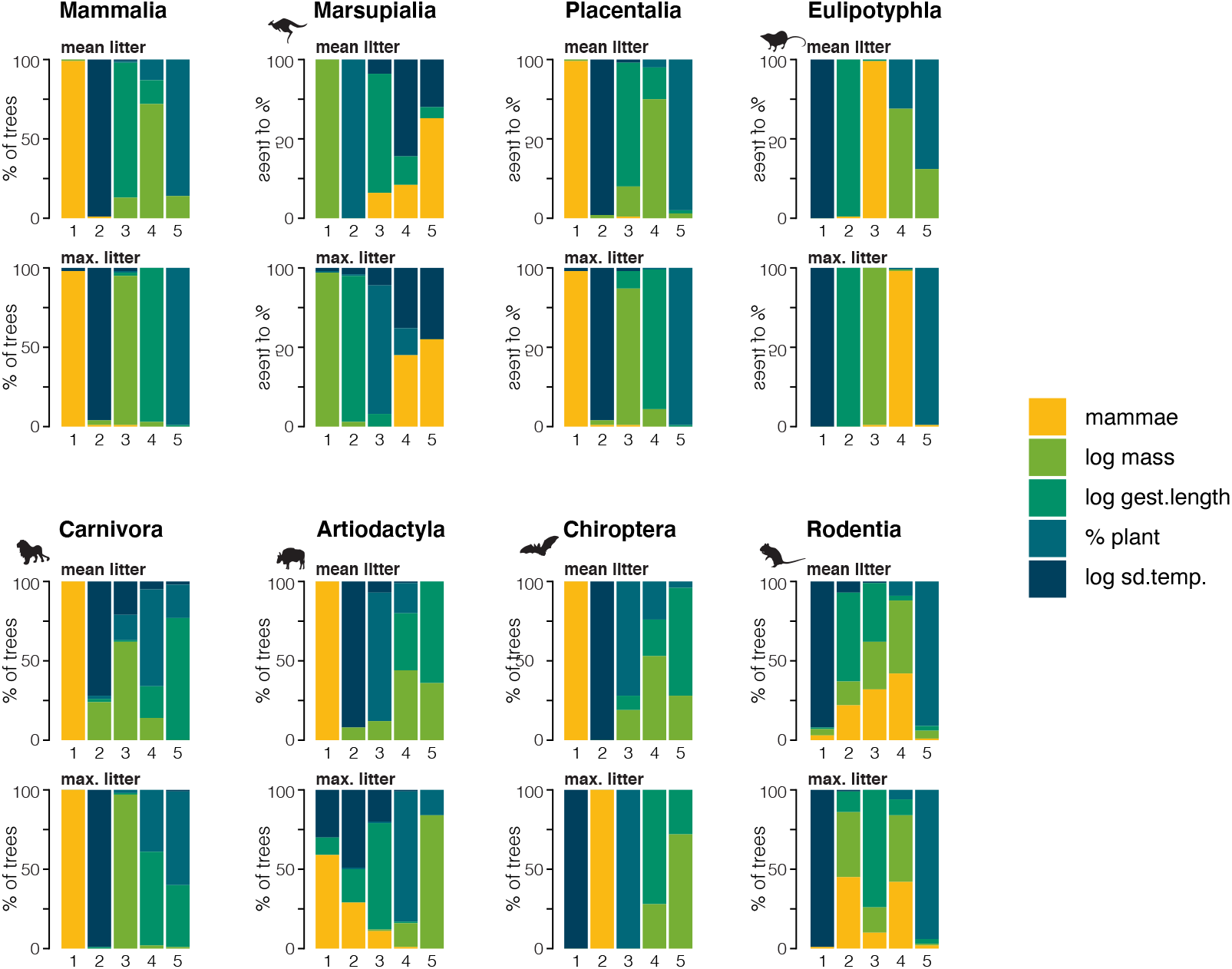
The relative predictive effect of five species-level traits on litter size. Likelihood testing was used to compare a five-predictor model with a series of models that sequentially excluded one of the predictor variables. Analyses were run for both mean and maximum litter size across 100 trees to account for phylogenetic uncertainty. The variables were ranked according to the magnitude of resultant ΔAIC for each of the 100 trees, with one being the greatest ΔAIC to five being the smallest ΔAIC. Stacked bar graphs show the percentage of trees where a variable occupied that particular rank. The first ranked variable of each clade corresponds with the results displayed in Fig. 3b.

**Supplementary Fig. 5.**
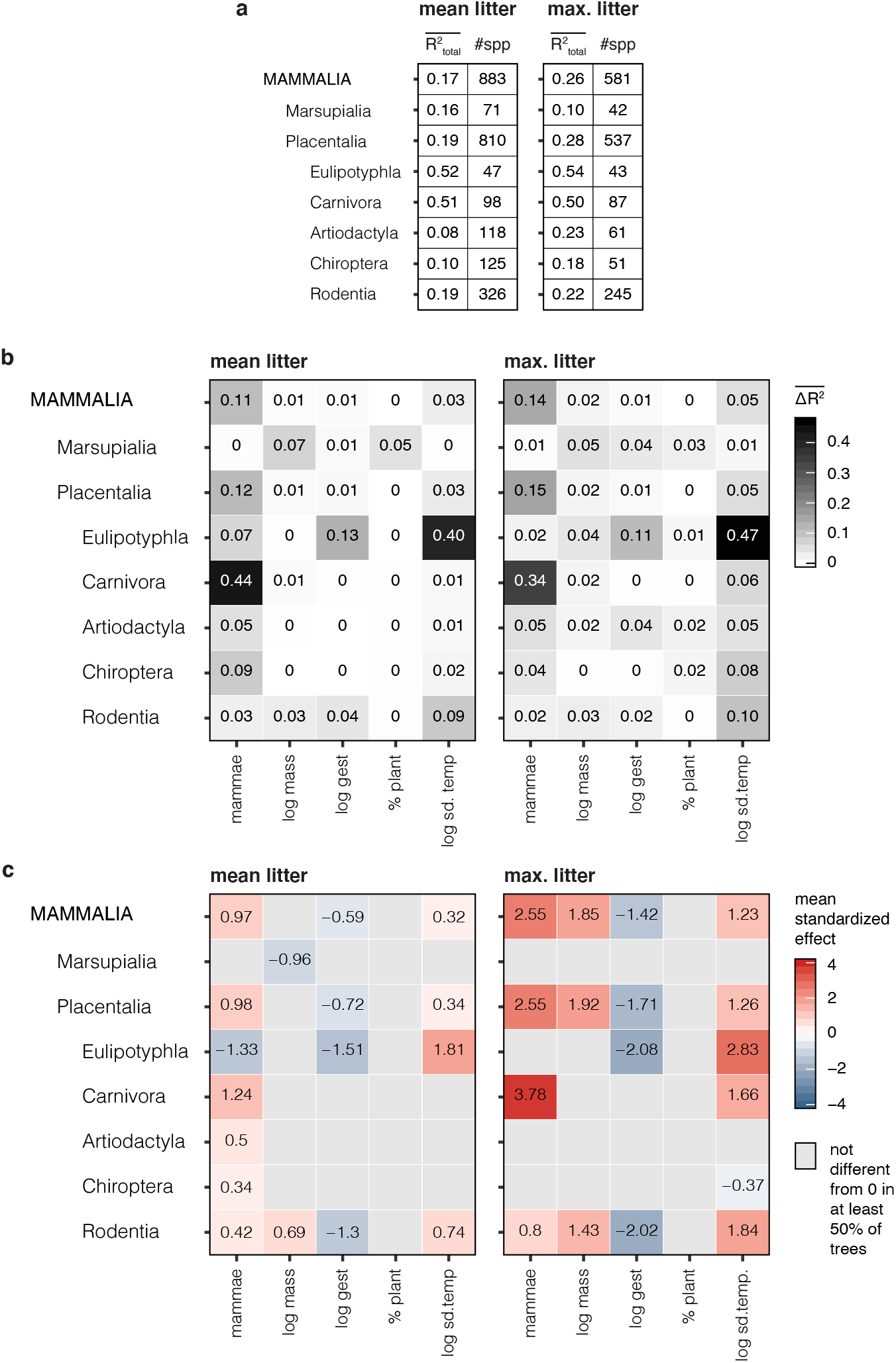
Additional results of analyses comparing five predictors on litter size. Likelihood testing was used to compare a five-predictor model with models that sequentially excluded one of the predictor variables. Analyses were run for both mean and maximum litter size across 100 trees to account for phylogenetic uncertainty, (a) Median R^2^ of the full five-parameter models and number of species analyzed, (b) The difference in R^2^ between the five-predictor model with the four-parameter models were calculated to evaluate the impact of each variable on total predictive effect. Heatmaps show the ΔR^2^ mean values calculated for 100 trees, (c) Standardized unique effects of each predictor variable were calculated from the full, five-predictor model. Heatmaps show the standardized mean effects calculated for the 100 trees. If standardized effects were not significantly different from zero to a threshold of *P* < 0.05 for at least 50% of the trees, the mean effect value is not shown, and the quadrant is represented by a grey box.

**Supplementary Fig. 6.**
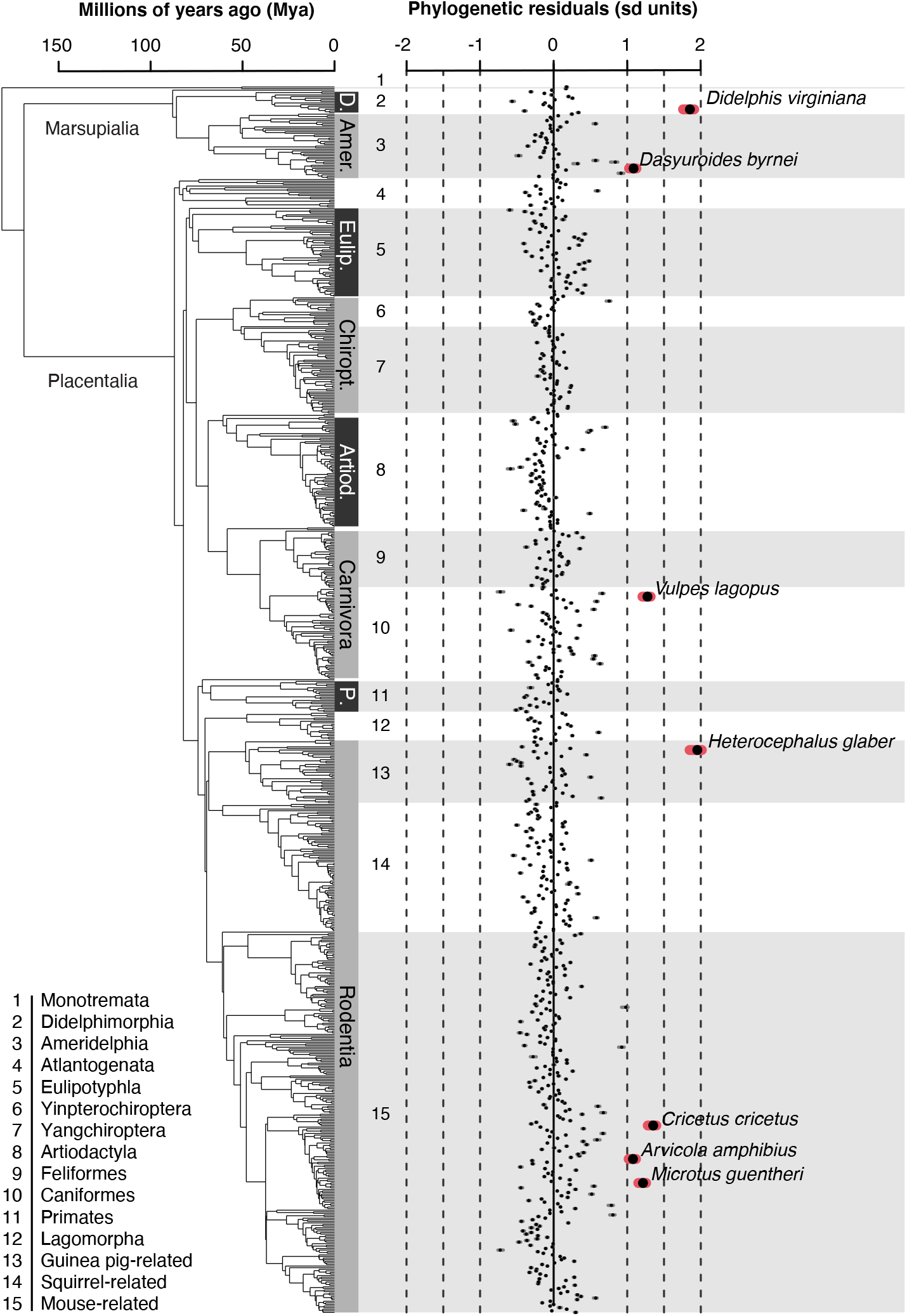
Phylogenetic residuals per species for the PGLS model of maximum litter size ~ mammae number. The residuals from the PGLS models run on each of 100 trees were multiplied by the factorized positive-definite matrix of phylogenetic variance-covariance, and then divided by the standard deviation to place in comparable units. Phylogenetic residuals were then plotted as median values (dots) and 95% confidence intervals (grey bars). Those species with phylogenetic residuals greater than 1 standard deviation are highlighted in red and labeled. Only the naked mole-rat (Heterocephalus glaber; maximum litter: 27, mammae: 6) and Virginia opossum (Didelphis virginiana; maximum litter: 32, mammae: 13) have phylogenetic residuals greater than 1.5. Abbreviations: D., Didelphimorphia; Amer., Ameridelphia; Eulip., Eulipotyphla; Chiropt., Chiroptera; Artiod., Artiodactyla; P. Primates.

**Supplementary Fig. 7.**
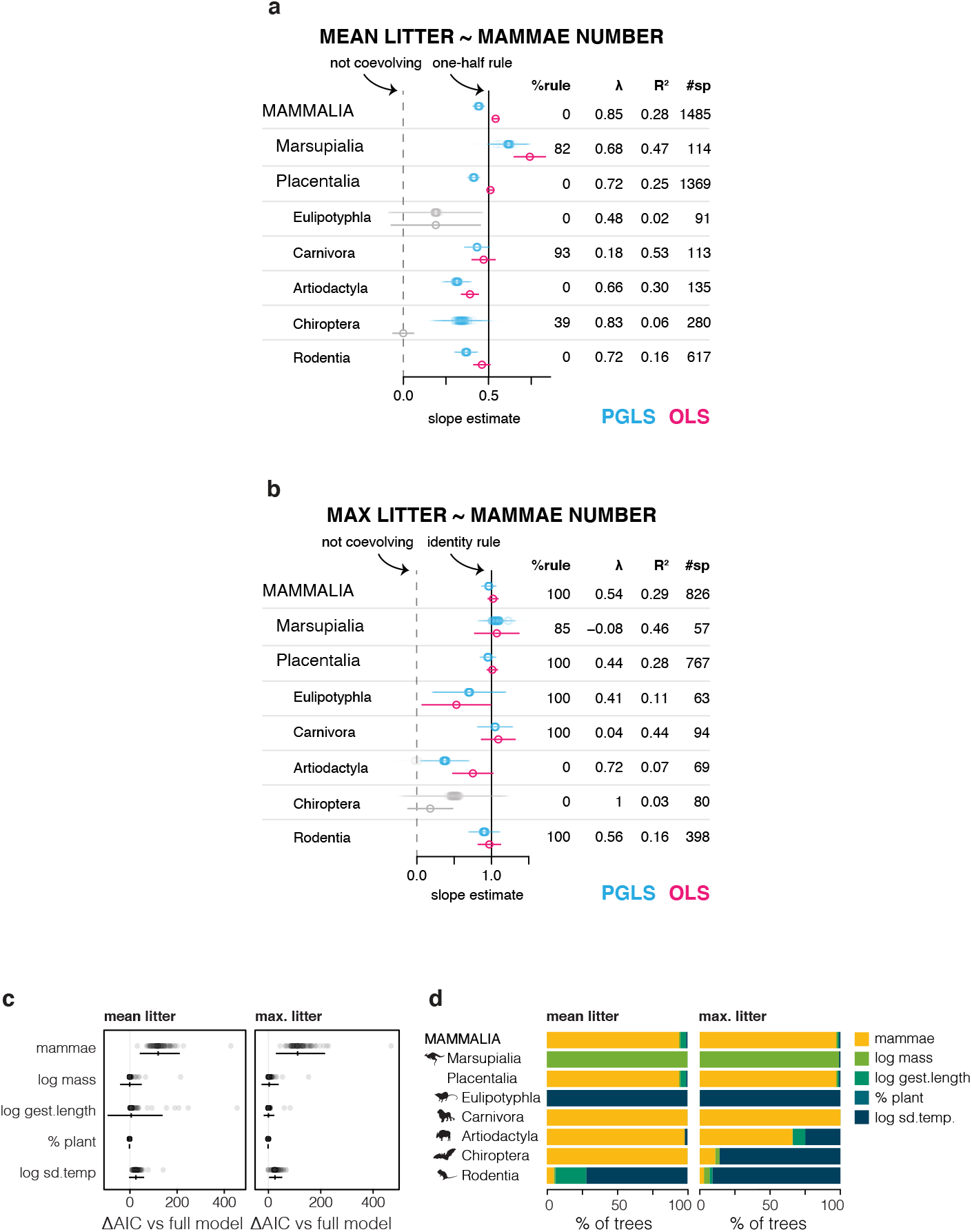
Testing sensitivity of PGLS results with the 5,911-species mammal species phylogeny. Analyses presented in the main body of the paper were implicated using an expanded species-level phylogeny of mammals and expanded dataset of mammae number, mean litter size, and maximum litter size, (a, b) PGLS and OLS regressions show consistent results to what is recovered by analyses of the 4,098-species time-calibrated molecular phylogeny, as shown in Fig. 2. (c, d) Multivariate regressions and likelihood model testing show consistent results to what is recovered by analysis of the 4,098-species time-calibrated molecular phylogeny, as shown in Fig. 3.

**Supplementary Fig. 8.**
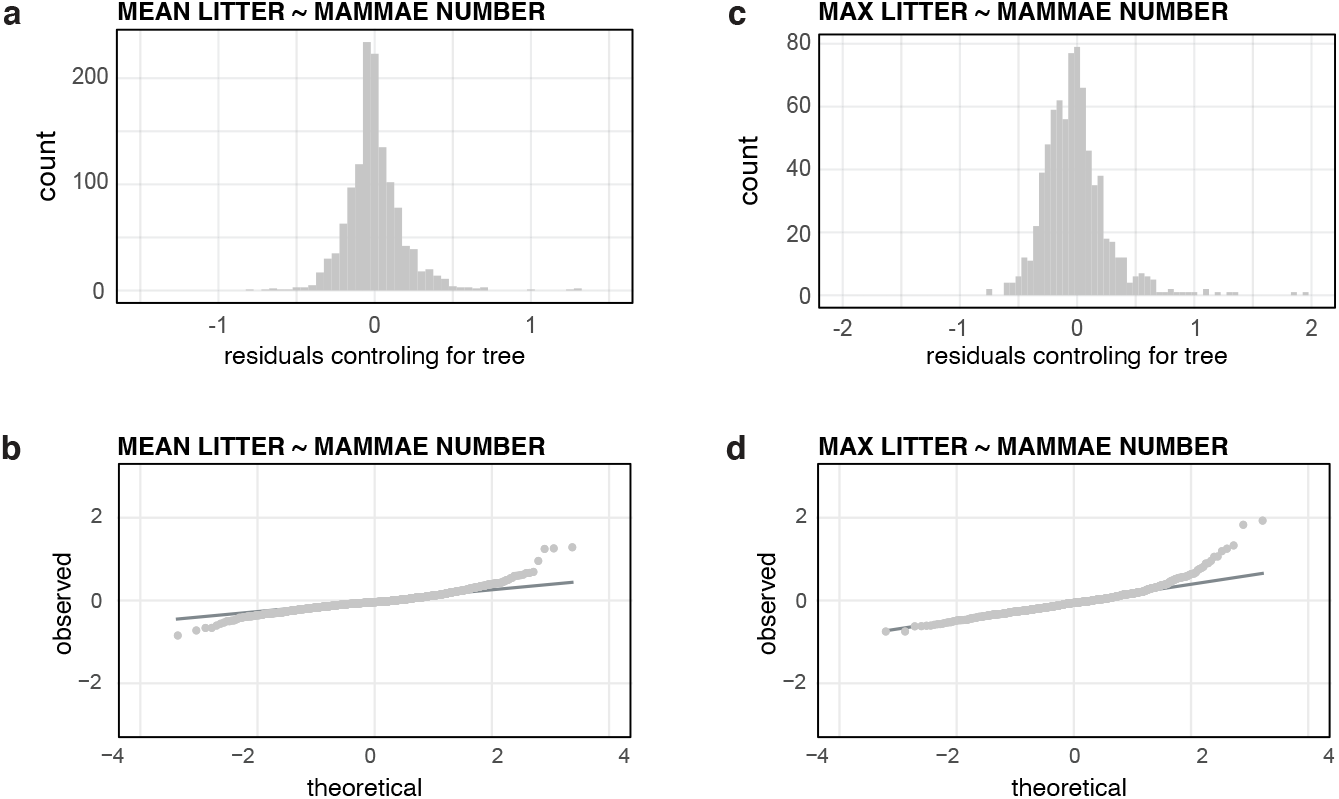
Visualizing the distributions of residuals from univariate PGLS regressions. The residuals from the PGLS models run on each of 100 trees were multiplied by the factorized positive-definite matrix of phylogenetic variance-covariance, and then divided by the standard deviation to place in comparable units, (a) Histogram of the median values of transformed residuals for each species in the PGLS regression of mean litter ~ mammae number, (b) Q-Q plot to assess normality of the median transformed residuals of PGLS regressions of mean litter ~ mammae number, (c) Histogram of the median values of transformed residuals for each species in the PGLS regression of maximum litter ~ mammae number, (d) Q-Q plot to assess normality of median transformed residuals of PGLS regressions of maximum litter ~ mammae. In neither regression do species show standardized residual values > 3.

**Supplementary Fig. 9.**
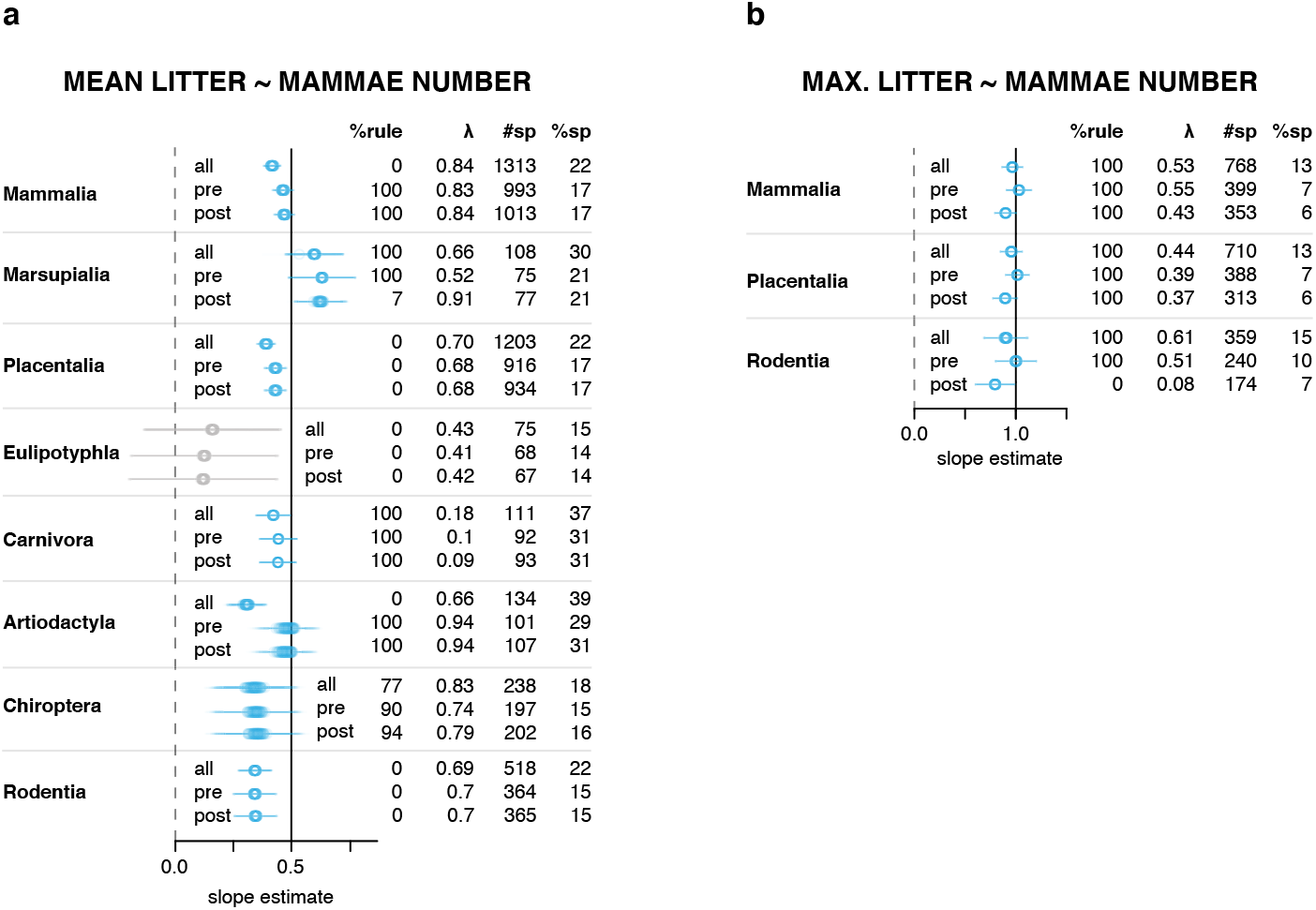
Testing sensitivity of PGLS analyses to timepoint of sampling. To assess how the time point of sampling could impact results, data subsets were generated that included only either pre-birth or post-birth observations, and PGLS regressions were run on Mammalia and on ordinal and infraclass clades for which data was available for at least 40 species, (a) PGLS regressions of mean litter ~ mammae number. In some regressions (e.g., Mammalia and Artiodactyla) estimates of the slope parameter differ between the full data set (labeled ‘all’) and the partitioned data sets (labeled ‘pre’ and ‘post’). We attribute this to reduced taxonomic coverage. However, results are consistent between the two timepoints in all cases, (b) PGLS regression of maximum litter ~ mammae number shows weak differences between pre- and post-birth data sets in Mammalia, Placentalia, and Rodentia, with slope estimates of the pre-birth data found to be greater than post-birth data estimates with the full data set as intermediate to these values. This pattern is suggestive of embryonic attrition and how sampling timepoint could impact parameter estimates. The result that Mammalia follows the identity rule is robust to such partitioning and, therefore, not sensitive to timepoint of embryonic observation. However, clades can be impacted by this effect (e.g., Rodentia).

**Supplementary Table 1 | Mammae number and litter size dataset.** Data on the mammae number, mean litter size, and maximum litter size as well as the sources for these observations. Taxonomic information is provided so that the data can be mapped to a phylogeny of mammals.

**Supplementary Table 2 | Tukey and Kramer test statistics.** Analyses were run to test for differences between clades in their distribution of mammae number, mean litter size and maximum litter size. Values presented are *P* values from the Tukey and Kramer tests, which correspond with Supplementary Fig. 2.

**Supplementary Table 3 | Summary of PGLS regressions run on 100 trees.** Phylogenetic generalized least squares regressions were run on 100 trees to test whether and how mammae number coevolves with mean litter size and maximum litter size. The summary table shows the percentage of regressions across 100 trees that were significant to the *P* value threshold of 0.05, the percentage that are significant and also have estimates of the slope and intercept parameters consistent with theoretical rules (‘one half rule’ and ‘identity rule’). We also report the mean estimate of Pagel’s λ (a measure of phylogenetic signal), the number of species included in the regression, and the percentage of extant species diversity that this reflects. These regressions are presented in Fig. 2, Supplementary Figs. 7, 9.

**Supplementary Table 4 | Summary statistics for the five-parameter additive model.** A five-parameter model was constructed to assess the explanatory power of mammae relative to other known correlates of litter size. Models were run over 100 trees and are summarized in the table. We report the number of species analyzed by each model and the mean R^2^ of the full five-parameter model. The impact of each predictor was calculated by dropping the variable and comparing the resultant 4-parameter model with the full model. For each predictor, we report mean ΔAIC, mean R^2^, mean standardized effect as well as the 95% confidence intervals for these results. Finally, for each of the 100 trees, the predictors were ranked according to magnitude of the ΔAIC they produced, and these rankings are summarized as a percentage of ‘ranked1’ through ‘ranked 6,’ which correspond from greatest to least ΔAIC. These analyses are presented in Fig. 3, Supplementary Figs. 4, 5.

**Supplementary Table 5 | Summary of OLS regressions.** Ordinary least squares regression were conducted for comparison with PGLS regressions. This summary table includes *P* values, adjusted R^2^ values, information pertaining to the *f* statistic, the number of species analyzed in the regression, the slope estimate and its standard error, the intercept estimate and its standard error, and whether the regression follows theoretical rules (‘one half rule’ and ‘identity rule’). These regressions are presented in Fig. 2 and Supplementary Fig. 7.

**Supplementary Table 6 | Ecological and life history trait dataset.** Data on the ecology and life history of mammal species that were used to construct the multivariate linear models.

